# Behavioral impairments are linked to neuroinflammation in mice with Cerebral Cavernous Malformation disease

**DOI:** 10.1101/2024.05.29.596485

**Authors:** Joseph Offenberger, Bianca Chen, Leigh-Ana Rossitto, Irisa Jin, Liam Conaboy, Helios Gallego-Gutierrez, Bliss Nelsen, Eduardo Frias-Anaya, David J. Gonzalez, Stephan Anagnostaras, Miguel Alejandro Lopez-Ramirez

## Abstract

**Background:** Cerebral Cavernous Malformations (CCMs) are neurovascular abnormalities in the central nervous system (CNS) caused by loss of function mutations in KRIT1 (CCM1), CCM2, or PDCD10 (CCM3) genes. One of the most common symptoms in CCM patients is associated with motor disability, weakness, seizures, stress, and anxiety, and the extent of the symptom or symptoms may be due to the location of the lesion within the CNS or whether multiple lesions are present. Previous studies have primarily focused on understanding the pathology of CCM using animal models. However, more research has yet to explore the potential impact of CCM lesions on behavioral deficits in animal models, including effects on short-term and long-term memory, motor coordination, and function.

**Methods:** We used the accelerating RotaRod test to assess motor and coordination deficits. We also used the open field test to assess locomotor activity and pathology-related behavior and Pavlovian fear conditioning to assess short—and long-term memory deficits. Our behavioral studies were complemented by proteomics, histology, immunofluorescence, and imaging techniques. We found that neuroinflammation is crucial in behavioral deficits in male and female mice with neurovascular CCM lesions (*Slco1c1-iCreERT2; Pdcd10^fl/fl^; Pdcd10^BECKO^*).

**Results:** Functional behavior tests in male and female *Pdcd10^BECKO^* mice revealed that CCM lesions cause sudden motor coordination deficits associated with the manifestation of profound neuroinflammatory lesions. Our findings indicate that maturation of CCM lesions in *Pdcd10^BECKO^* mice also experienced a significant change in short- and long-term memory compared to their littermate controls, *Pdcd10^fl/fl^* mice. Proteomic experiments reveal that as CCM lesions mature, there is an increase in pathways associated with inflammation, coagulation, and angiogenesis, and a decrease in pathways associated with learning and plasticity. Therefore, our study shows that *Pdcd10^BECKO^* mice display a wide range of behavioral deficits due to significant lesion formation in their central nervous system and that signaling pathways associated with neuroinflammation and learning impact behavioral outcomes.

**Conclusions:** Our study found that CCM animal models exhibited behavioral impairments such as decreased motor coordination and amnesia. These impairments were associated with the maturation of CCM lesions that displayed a neuroinflammatory pattern.

## Introduction

Cerebral cavernous malformations (CCMs) are common neurovascular lesions that cause a lifelong risk of brain hemorrhage, thrombosis, seizures, and neurologic deficits. Currently, no effective pharmacologic treatment is available ^1–5^. At present, only a few symptoms can be relieved through drug treatments, such as managing headaches and seizures. Surgery is frequently not feasible for lesions located in critical or hard-to-reach places or when patients have multiple lesions within the central nervous system (CNS). An estimated 0.2% of the US population has a CCM lesion, but only 25% are diagnosed^6, 7^. Of these, ∼20% are diagnosed as children (0.0125% of US population or approximately 40,000 patients)^8^. Around 20% of CCM cases are familial. Familial CCM is an autosomal dominant disease with variable penetrance, even within the same family^9^. Familial cases usually exhibit multiple lesions, whereas sporadic cases (approximately 80% of CCM cases) of CCM usually exhibit an isolated lesion. The symptoms most frequently found in patients with CCMs are related to motor disability, weakness, seizures, stress, and anxiety^10^. The severity of these symptoms may depend on the location of the lesion within CNS, the stage of the lesions (active vs inactive), or the presence of multiple lesions. There is a variable impact on life expectancy, with mortality resulting from uncontrolled seizure and hemorrhage or surgical sequelae in some cases.

Previous studies have proposed a vascular hypothesis that maintains that CCMs are initiated by vascular endothelial cell dysfunction by loss of function mutations (which can be inherited germline and somatic mutations) in the genes *KRIT1* (Krev1 interaction trapped gene 1, *CCM1*), *CCM2* (*Malcavernin*) or *PDCD10* (Programmed cell death protein 10, *CCM3*)^11–14^ propel brain vascular changes (*<u>CCM endothelium</u>*), marked by the disruption of intercellular junctions^15–18^, increase in angiogenesis^16, 19–22^, endothelial-mesenchymal transition^23^, reactive oxygen species^11, 24–26^, vascular permeability^15, 27–38^ and hypoxia signaling^4, 39^. Human and mouse studies have also revealed that inflammation contributes to brain vascular malformations ^3, 4, 40–43^. Human CCM lesions tend to bleed, which may lead to inflammation and thrombosis in varying degrees^44–46^. Recent findings in CCM animal models further support that crosstalk between astrocytes and CCM endothelium^3, 4^ can trigger the recruitment of inflammatory cells arising from brain parenchyma (microglia) and periphery immune system (leukocytes) into mature CCM lesions that propagate lesion growth, thrombosis, and bleeding, which contribute to heterogeneity in CCM severity ^3, 40, 47,48^.

Our research revealed that male and female mice with *Pdcd10* gene knockout in brain endothelial cells (*Pdcd10^BECKO^*) exhibit sudden motor coordination deficits associated with neuroinflammatory CCM lesions. We found that *Pdcd10^BECKO^* mice also display significant deficits in short- and long-term memory compared to their littermate controls, *Pdcd10^fl/fl^* mice. Our study further reveals that *Pdcd10^BECKO^*mice show extensive behavioral deficits resulting from significant lesion formation, primarily in the hippocampal region. We observed that activating signaling pathways associated with neuroinflammation, thrombosis, and learning affect the maturation of CCM lesions that correlate with behavioral deficits. Therefore, our study proposed that animal models of CCM exhibit behavioral impairments, including impaired motor coordination and amnesia, due to neuroinflammation-driven maturation of CCM lesions.

## Material and Methods

### Genetically modified animals

Brain endothelial-specific conditional *Pdcd10*-null mice were created by crossing a *Slco1c1* promoter-driven tamoxifen-regulated Cre recombinase (*Slco1c1-CreERT2,* a gift from Markus Schwaninger, University of Lubeck) strain with LoxP-flanked *Pdcd10* (*Pdcd10^fl/fl^*, a gift from Wang Min, Yale University; *Slco1c1-CreERT2; Pdcd10^fl/fl^*) mice. On a postnatal day 5 (P5), mice were administered 50 μg of 4-hydroxy-tamoxifen (H7904, Sigma-Aldrich) by intragastric injection to induce genetic inactivation of the endothelial *Pdcd10* gene in littermates with *Slco1c1-CreERT2;Pdcd10^fl/fl^* (*Pdcd10^BECKO^*), and *Pdcd10^fl/fl^* mice were used as littermate controls. We also used non-injected *Slco1c1-CreERT2; Pdcd10^fl/fl^* mice as littermate controls whose gene expression and histology were the same as in *Pdcd10^fl/fl^* mice. All animal experiments complied with animal procedure protocols approved by the University of California, San Diego’s Institutional Animal Care and Use Committee and compliant with the 8^th^ *Guide for Care and Use of Laboratory Animals*.

### Neurobehavioral test

Male and female mice were used for behavior outcomes. Animals were group-housed (2-5 animals per same-sex cage), given unrestricted access to food and water, and kept on a 14:10-h light/dark schedule; all behavioral testing occurred during the light phase. Mice were at least 4 weeks old and handled for 3 days (1 min/day) before testing.

### Rotarod performing test

Effects on motor coordination were assessed using a RotaRod performance test. The RotaRod apparatus consists of 5 individual slots containing a rotating horizontal dowel upon which mice are placed (ENV-575M, Med-Associates Inc., Georgia, VT). Mouse slot positions on the RotaRod were counterbalanced across testing to control for any variability in the chamber environment. The RotaRod procedure entailed two phases. First mice underwent a training phase (30s at 3 rpm) in order to learn the RotaRod task. This was followed by an acceleration phase (3 - 30 rpm over 300 s). Training and acceleration phases were performed in light conditions. A 30-min resting period was implemented between each acceleration run during the first day (training), in which mice were returned to their cages with access to food and water. Further testing was performed once a week for three weeks. Motion sensors detected the fall of each mouse from the dowel, ending the test for that animal and recording the latency to fall from the start of the test^1^.

### Fear conditioning test

The VideoFreeze system (Med-Associates Inc., Georgia, VT) was used to test freezing behavior and locomotor activity via digital video^49^. Freezing is an index of fear memory, as mice will freeze in response to conditioned fear stimuli. Testing took place in individual conditioning chambers housed in a windowless room. Each chamber (32 X 25 X 25 cm) was located within an outer, sound-attenuating chamber (63.5 X 35.5 X 76 cm; Med-Associates Inc., Georgia, VT) and equipped with a speaker in the side wall, a stainless steel grid floor (36 rods, each rod 2-mm diameter, 8-mm center to center; Med-Associates Inc., Georgia, VT), and stainless steel drop pan. Each testing chamber was equipped with an overhead LED light source providing both visible broad-spectrum white light (450–650 nm) and near-infrared light (NIR; 940 nm), and an IEEE 1394 progressive scan video camera with a visible light filter (VID-CAM-MONO-2A; Med-Associates Inc., Georgia, VT) connected to a computer and video equipment in an adjacent room. Each testing chamber was also connected to a solid-state scrambler, providing AC constant current shock and an audio stimulus generator, controlled via an interface connected to a Windows computer running the VideoFreeze software.

For fear conditioning, all 32 mice were transported from their home cages in an adjacent room and placed in the conditioning chambers. After recording 2 min of baseline activity, a 30 s tone (2.8 kHz, 85dBA) was presented and co-terminated with a scrambled footshock (2 s, 0.75mA, AC constant current) delivered through the rod floor of the cage. The training trial continued for 2.5 min after the termination of the footshock, followed by an additional 5-minute test of immediate memory (post-shock freezing) in the same chamber. Before each trial, chambers were cleaned and scented with 7% isopropyl alcohol to provide a background odor. Background white noise (65-dBA) was provided by a speaker located centrally in the room. Following each trial, mice were returned to their home cages.

### Context memory test

One week after fear conditioning, mice were returned to the training chambers for an assessment of contextual fear memory. During this assessment, no shock was delivered and all contextual cues were identical to those provided during the fear conditioning phase. Freezing to the fear conditioning context was scored over a 5-minute testing period, and mice were then returned to their home cages.

### Tone memory test

Twenty-four hours after the context memory test, mice were given a tone memory test to assess cued fear memory. For testing cued fear memory, the conditioning context was modified along several dimensions. Black acrylic sheets were placed over the grid floor to provide an alternate sensory experience. Black plastic triangular tents, translucent only to NIR light, were placed inside each box, with each side of the triangle measuring 23 cm. Only NIR light was used, creating a completely dark environment visible only to the video camera. Before each trial, chambers were cleaned and scented with a 5% acetic acid solution to provide a background odor, and background white noise (65-dB) was provided by a speaker located centrally in the room. Baseline activity was assessed for 2 min, after which a 30s tone (identical to the training tone) was presented at minutes 2, 3, and 4. After the tone test, the mice were returned to their home cages. During each phase, freezing was measured by a computer running VideoFreeze.

### Open field test

Mice underwent open field testing (OFT) to assess their activity without aversive or appetitive stimuli for 30 min. For OFT, seven mice were tested concurrently in individual open-field chambers (Med Associates Inc., St. Albans, VT, USA). Each chamber (21.5 X 21.5 X 31 cm) had the same visual and tactile properties (stainless steel rod floors with clear polycarbonate walls), and mice were tested in moderate light (∼15 lx). Before testing between trials, each chamber was cleaned with 10% glass cleaner (Zep Inc., Atlanta, GA, USA). A central speaker in the room supplied background white noise (65dBA). Experimenters placed individual animals in the center of one of seven open field chambers and left the room. Activity Monitor software (Med Associates Inc.) used the interruption of infrared beams in *x, y,* and *z* planes to identify mouse position and score locomotion (distance), average ambulatory speed (cm/s), resting time (s), stereotypy (counts), and verticality (counts). Stereotypy is persistent, repetitive behavior for no apparent purpose, measured by rapid breaks of the same photobeam. Verticality is the animal’s vertical rearing in the testing apparatus and is counted when the animal breaks the top-most beam in the *z* plane. At the end of the 30-minute trial, animals were returned to their home cages^2, 3^.

### Histology

Brains from *Pdcd10^BECKO^* mice were isolated and fixed in 4% paraformaldehyde (PFA) in PBS at 4°C overnight. Tissue was cryoprotected in 30% sucrose solution in PBS and then embedded and frozen in an O.C.T compound (Fischer Scientific). Brains were cut using a cryostat into 18-µm sagittal sections onto Superfrost Plus slides (VWR International). Sections were stained by hematoxylin and eosin. The slides were viewed with a high-resolution slide scanner (Olympus VS200 Slide Scanner), and the images were captured with VS200 ASW V3.3 software (Olympus).

### Quantitative Multiplex Proteomics

Peptide preparation: Whole brain lysates were made by homogenizing flash-frozen cortical dissections in lysis buffer (6M urea, 7% SDS, 50 mM TEAB, titrated to pH 8.1 with phosphoric acid) with protease and phosphatase inhibitors added (Roche, CO-RO and PHOSS-RO) and then sonicated. Disulfide bonds were reduced in 5 mM dithiothreitol (DTT) at 47°C for 30 min, and free cysteines were alkylated in 15 mM iodoacetamide in a room temperature, darkened environment for 20 min. The alkylation reaction was quenched for 15 minutes at room temperature by adding an equivalent volume of DTT as in the reduction reaction. Proteins were then trapped using ProtiFi S-Trap columns (ProtiFi, C02-mini), digested with sequencing-grade trypsin (Promega, V5113), and eluted according to manufacturing protocols. Eluents were desalted on SepPak C18 columns using instructions provided by the manufacturer (Waters, WAT054960). Desalted samples were dried under a vacuum. Peptides were quantified using a Pierce Colorimetric Peptide Quantification Assay Kit following the manufacturer’s instructions (Thermo Scientific, 23275). 50 μg of each sample was separated for multiplex proteomic analysis, with several samples aliquoted twice to serve as technical duplicates and fill multiplex channels. Two identical bridge channels, 50 μg each, were created by pooling equal amounts of each sample.

Labeling: This experiment required two Tandem Mass Tag (TMT) 16plexes (Thermo Scientific, A44520; lot number WJ327115). The labeling schemes for multiplex experiment are included in the Supplemental Material. Sample duplicates to fill empty channels were randomly chosen, bridge channels were assigned to channel 126, and the samples were randomly assigned to the remaining channels across the two plexes. Before labeling, dried peptides were resuspended in 50 μL of 30% acetonitrile (ACN) with 200 mM HEPES, pH 8.5, and TMT labels were suspended in anhydrous ACN to a final concentration of 20 mg/mL. Each resuspended sample was mixed with 7 μL of the appropriate tag, and the labeling reaction proceeded at room temperature for one hour. Excess label was quenched with 8 μL of 5% hydroxylamine for 15 minutes. 50 μL of 1% trifluoroacetic acid (TFA) was added to each sample, then samples were combined into the appropriate multiplexes. Plexes were desalted on SepPak C18 columns and dried under a vacuum.

Fractionation: The two 16plexes were next subjected to fractionation using reverse phase high pH liquid chromatography to increase sequencing depth. An Ultimate 3000 high-performance liquid chromatography system fitted with a fraction collector, C18 column (4.6 x 250 mm), solvent degasser, and variable wavelength detector was used for fractionation. Plexes were resuspended in 105 μL of 25 mM ammonium bicarbonate (ABC) and were fractionated on a 10-40% ACN gradient with 10 mM ABC for over an hour^50^. The resulting 96 fractions were concatenated into 24 fractions by combining alternating wells within each column, and 12 alternating fractions were used for mass spectrometry analysis, as previously described^51^.

Mass spectrometry-based proteomic analysis: Fractions were resuspended in 5% FA/5% ACN at a concentration of 1 μg/μL and analyzed on an Orbitrap Fusion Tribrid mass spectrometer with an in-line Easy-nLC 1000 System. 1 μg of each fraction or sample was loaded onto a 30-cm in-house pulled and packed glass capillary column (I.D. 100 μm, O.D. 350 μm). The column was packed with 0.5 cm of 5 μm C4 resin followed by 0.5 cm of 3 μm C18 resin, with the remainder of the column packed with 1.8 μm of C18 resin. Electrospray ionization was assisted by the application of 2000 V of electricity through a T-junction connecting the column to the nLC, heating the column to 60°C. All data acquired were centroided.

Peptides were eluted after loading using a gradient ranging from 6% to 25% ACN with 0.125% formic acid over 165 minutes at a flow rate of 300 nL/min. Data were acquired in data-dependent mode with polarity set to positive. Dynamic exclusion was set to exclude after n=1 times for 90 seconds. Charge states 2-6 were included. Maximum injection time was set to 100 ms. MS1 spectra were acquired in the Orbitrap with a scan range of 500-1200 m/z and a mass resolution of 120,000. Ions selected for MS2 analysis were isolated in the quadrupole with a 0.7 m/z isolation window. Ions between 600-1200 m/z and a minimum intensity of 5e3 were fragmented using collision induced dissociation, with a normalized collision energy of 30% and activation time of 10 ms. Ions were detected in the ion trap with a rapid scan rate and a maximum injection time of 50 ms. MS3 analysis was conducted on ions between 400-2000 m/z using the synchronous precursor selection option to maximize TMT quantitation sensitivity. MS2 ions 40 m/z below and 15 m/z above the MS1 precursor ion were excluded from MS3 selection. Up to 10 MS2 ions were simultaneously isolated and fragmented with high-energy collision–induced dissociation using a normalized energy of 50% and a maximum ion injection time of 250 ms. MS3 fragment ions were analyzed in the Orbitrap at a resolution of 60,000.

Data processing and normalization: Raw mass spectrometry files were processed using Proteome Discoverer 2.5. Using the SEQUEST algorithm, MS2 data were queried against Uniprot *Mus musculus* (mouse) reference proteome UP000000589. A decoy search was also conducted with sequences in reverse order. Data were searched using a precursor mass tolerance of 50 ppm and fragment mass tolerance of 0.6. Static modifications were specified as follows: carbamidomethylation of cysteines and TMT reagents on lysines and N-termini as appropriate. Dynamic modifications were specified to include oxidation of methionines. The enzyme specificity was set to full trypsin digest with two missed cleavages permitted. Resulting peptide spectral matches were filtered at a 0.01 FDR by the Percolator module against the decoy database. Peptide spectral matches were quantified using precursor ion or MS3 ion intensities for label-free and labeled experiments, respectively.

Peptide spectral matches were exported and summed to the protein level, using only spectra with a signal-to-noise per label >10 and isolation interference of <25%. Plexes were then batch corrected in a multi-step process, as previously described. Briefly, within each plex, data are standardized to the average for each protein and then to the median of all averages for protein. Then, to account for slight differences in amounts of protein labeled, these values are then standardized to the median of the entire plex dataset^52^. Following batch correction, each sample transformed using Box-Cox power transformation (*MASS* R package version 7.3-55), abundances were scaled between 0-1 (*reshape* R package version 0.8.8), and each sample was standardized by its mean^52^. Technical duplicates were averaged at this step. Final data represent normalized relative protein abundances.

### Statistical analysis

Statistical analysis was performed using Prism software (GraphPad). Data are expressed as mean values ± standard error of the mean (SEM) for multiple individual and biological experiments, unless otherwise specified. The number of independent and biological replicates (n) is indicated for all experiments. Mice experiments were randomized, and histological quantification was performed blinded. The Shapiro-Wilk normality test with alpha=0.05 assessed the normality of data. For comparison between two groups, Student’s unpaired two-tail *t-test* or analysis of variance (ANOVA) for comparison among multiple groups, followed by the Tukey post hoc test, was used for normally distributed data. Mann-Whitney two-tailed test was used for two groups, and the Kruskal-Wallis test, followed by Dunn’s post hoc test, was used for non-normally distributed data. Assuming means and standard deviation from previous studies, we calculated sample sizes using a two-sided alpha of 0.05 and 80% power while assuming homogeneous variances for two samples^4,13,15,16^. At least three biologically independent experiments were conducted to ensure reproducibility. For proteomics: Principal component analysis of normalized protein abundances was performed using prcomp (R package *stats* version 4.4.0) and visualized using autoplot (R packages *ggplot* version 3.5.1, *ggfortify* version 0.4.13). Binary comparisons were performed on *Pdcd10*^BEKO^ against age-matched *Pdcd10*^fl/fl^ controls. Significantly differentially abundant proteins were assessed by a combination of Welch’s t-tests, which assumes normalized data with equal means and unequal variances, with a p-value of <

0.05 and a π-score (-logP•|log_2_(fold-change)|) of > 1^53^. Overlapping differentially abundant proteins were visualized with UpSet plots (R package *UpSetR* version 1.4). Gene ontology analysis on overlaps was performed using MetaScape (v3.5.20240101)^54^. RNA sequencing data ^48^ was integrated into the proteomics analysis using P60 mice from the current study and P55 normoxia mice from the prior study, and overlapping genes (n=5706) were analyzed. Correlation analysis was performed in GraphPad Prism using simple linear regression and Spearman’s r. R-generated graphs were edited in Adobe Illustrator for aesthetics.

## Data availability

R Proteomic data and additional supplementary files for reanalyzing the data collected here are available online at https://massive.ucsd.edu

## Results

### Pdcd10^BECKO^ mice exhibit impairments in motor coordination and balance

Patients with CCM disease commonly experience symptoms such as motor disability, weakness, seizures, stress, and anxiety. The severity and type of symptoms may vary depending on the location, number of lesions, and lesion activity in the CNS. Therefore, we first aimed to determine whether there is a correlation between the aggravation of CCM lesions and behavioral deficits in CCM animal models. We generated the *Pdcd10^BECKO^* (brain microvascular endothelial cell-specific *Pdcd10* knock-out) animal model by crossing the brain endothelial tamoxifen-regulated Cre (*Slco1c1-CreERT2*) with *Pdcd10^fl/fl^* mice (*Slco1c1-CreERT2; Pdcd10^fl/fl^*)^4, 39^ (Fig. 1A). Upon tamoxifen injection, *Pdcd10^BECKO^* animals develop progressive CCM lesions throughout their brains and neuroinflammation play a critical role in transitioning from initial CCM formation to mature and active lesions (aggravation of CCM lesions)^39^. We utilized the well-established accelerating RotaRod test to assess motor behavioral outcomes in CCM mice models. This test is particularly sensitive to motor function, coordination, and balance deficits. We initially observed no significant difference in the performance of *Pdcd10^BECKO^* and control *Pdcd10^fl/fl^* mice on the RotaRod test at postnatal days 35 (P35). This indicates that motor coordination was comparable between groups at this stage (Fig. 1B). Notably, after P42, we observed that *Pdcd10^BECKO^*mice showed decreased motor coordination, as evidenced by lower performance on the RotaRod test (Fig. 1A) and extensive cerebral lesion burden (Fig. 2). Furthermore, the lower performance on the RotaRod test persisted in *Pdcd10^BECKO^* mice until the end of the experimental period at P56 (Fig. 1B). These results suggest that neuroinflammation in CCM lesions may significantly contribute to the disease’s disability, as the inflammation pathway initiates approximately at P40 in this CCM animal model^39^. We conducted an open field test to evaluate the locomotor activity and pathology-related behaviors of P49 *Pdcd10^BECKO^* mice. Our observation showed that these mice exhibit a significant decrease in the level stereotypy, indicating potential hypoactivity, behavioral dissimilarities to psychiatric disorders with stereotypical-perseverative symptoms, and/or decreased anxiety-like behavior associated with CCM lesions (Supplemental Fig. 1B). Stereotypy is commonly associated with a variety of psychiatric illnesses and may indicate a behavioral target of future investigation in this model. These mice also exhibited increased average resting time(s), possibly indicating increased lethargy, hypolocomotion, and/or pain sensitivity (Supplemental Fig 1C). However, we did not observe any significant differences in verticality (counts), total movement distance (measured in cm) or speed (measured in cm/s) in *Pdcd10^BECKO^* mice and the control group *Pdcd10^fl/fl^* (Supplemental Fig 1).

**Figure 1.**
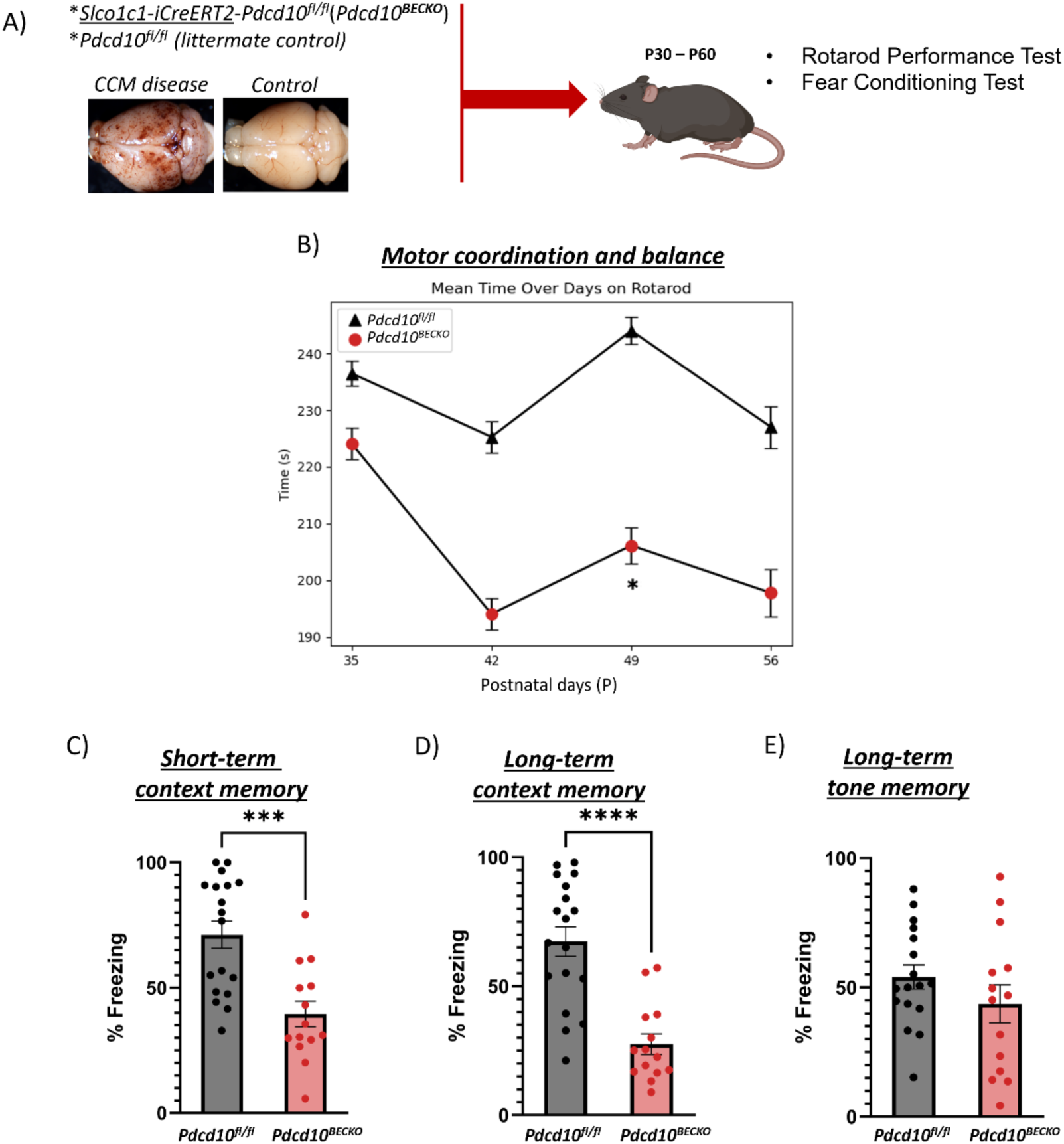
*Pdcd10^BECKO^* mice exhibit neurobehavioral deficits. A) A brain endothelial-specific CCM animal model (*Slco1c1-CreERT2; Pdcd10^fl/fl^*[*Pdcd10^BECKO^*], and control *Pdcd10^fl/fl^* mice) was employed for motor, open field and learning and memory performance. B) *Pdcd10^BECKO^* male and female mice show no significant differences at P35 (n = 15 *Pdcd10^fl/fl^*and 17 *Pdcd10^BECKO^*, *P* = 0.558), but acute motor deficits using the RotaRod test beginning at P42 (n = 18 *Pdcd10^fl/fl^*and 14 *Pdcd10^BECKO^*) throughout the end of the experimental period. Data are presented as a mean ± SEM. C) *Pdcd10^BECKO^* male and female mice show contextual learning and memory deficits using the fear conditioning test at P50-P55. D) *Pdcd10^BECKO^* male and female mice show contextual learning and memory deficits using the fear conditioning test at P57-P63 (n = 18 *Pdcd10^fl/fl^* and 14 *Pdcd10^BECKO^*). E) *Pdcd10^BECKO^* male and female mice show no tone learning and memory deficits using the fear conditioning test at P57-P63. Data are presented as a mean ± SEM. \**P* < 0.05, \*\**P* < 0.01, \*\*\**P* < 0.001.

**Figure 2.**
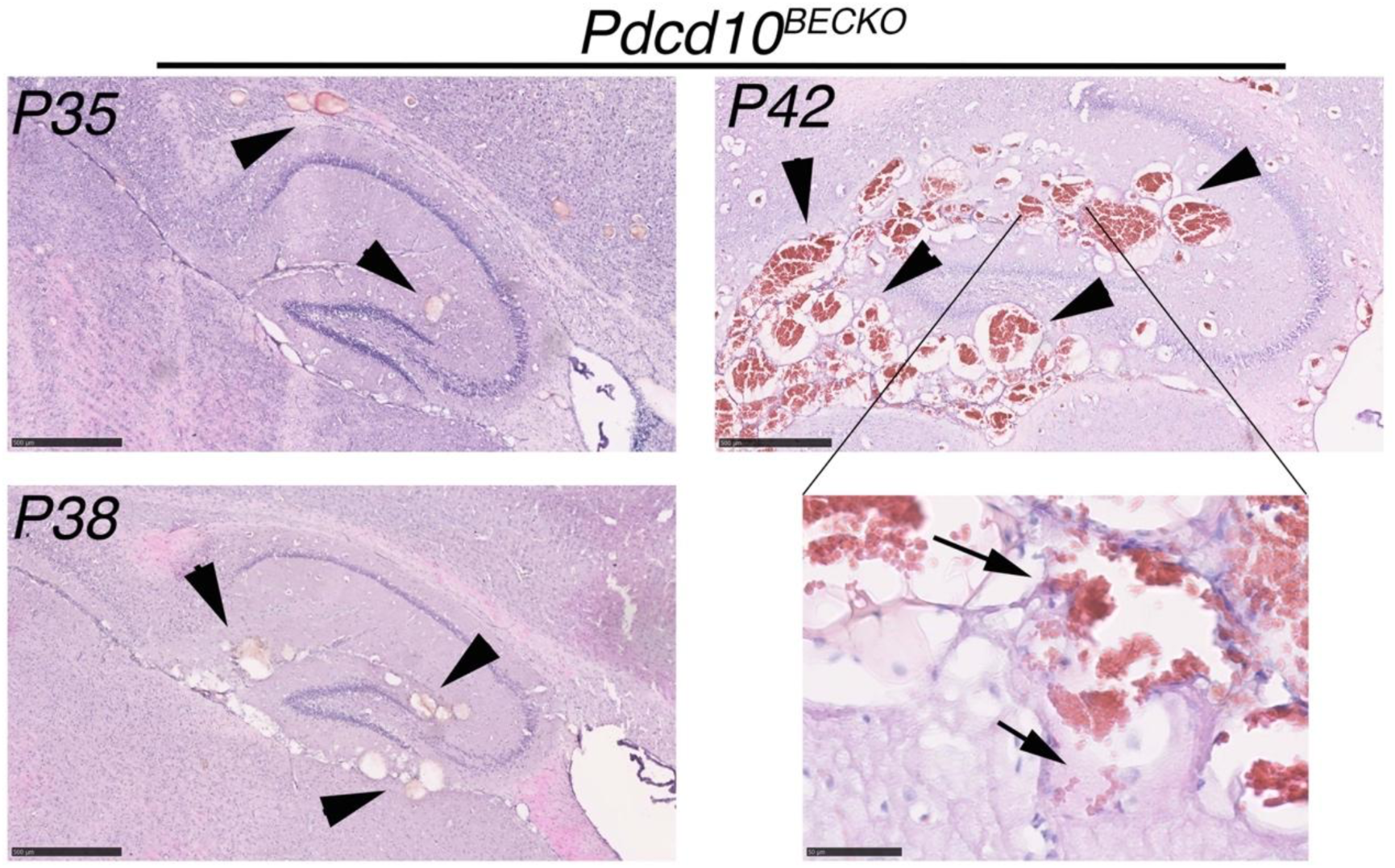
*Pdcd10^BECKO^* mice display behavioral deficits linked to an increase in CCM lesion burden. Histological analysis of brain sections from P35, P38, and P42 *Pdcd10^ECKO^* mice. Multiple CCM lesions were detected in sections stained with hematoxylin and eosin (Pink) at the hippocampal brain region. The arrowhead indicates CCM lesions, while the arrows show intra and extra-lesional bleeding present in P42 *Pdcd10^BECKO^* brains. Scale bars are 500 μm and 50 μm for high magnification.

### Pdcd10^BECKO^ mice exhibit amnesia specific to hippocampus-dependent memory

We investigated the effects of neurovascular CCM lesions on short- and long-term memory using Pavlovian fear conditioning. In this test, the mice experience an auditory cue accompanied by an electric foot shock, and freezing behavior is assessed during a post-shock immediate memory test and one week later during contextual and cued fear memory tests to evaluate short and long-term fear memory. Contextual memory is of particular interest because it is hippocampus-dependent and considered a model of human memory for facts and events^4–6^ (i.e., declarative). Therefore, increased freezing time is associated with better learning and memory. We observed that even when P56 *Pdcd10^BECKO^* mice had extensive CCM lesions in their brain and spinal cord^39^, they exhibited similar locomotor activity to the *Pdcd10^fl/fl^* mice control during the training baseline period (Supplemental Fig. 2). Additionally, a single tone-shock (Shock 1) pairing elicited an unconditioned response in P56 *Pdcd10^BECKO^*and *Pdcd10^fl/fl^* mice to the same extent (Supplemental Fig. 2). This indicates that the baseline and the response to stimuli before training were comparable for both groups at this stage. We collected additional data on the same animals to quantify learning and memory differences between *Pdcd10^BECKO^* mice and *Pdcd10^fl/fl^* controls. First, we noticed significant behavioral deficits in short-term memory in P63 *Pdcd10^BECKO^* mice compared to the control *Pdcd10^fl/fl^* mice (Figure 1C). Additionally, long-term cognitive deficits^7^ can be induced in P63 *Pdcd10^BECKO^* mice (Figure 1D), which have been observed to experience extensive neuroinflammation^39, 48^. No significant group differences were observed for long-term context tone memory (Figure 1E). Therefore, these results indicate that *Pdcd10^BECKO^* mice exhibit significant impairment in both short-term and long-term contextual memory.

### Whole-brain proteome profiling reveals increased neuroinflammation and decreased learning and plasticity proteins in Pdcd10^BECKO^ mice

During CCM disease, we and others were able to show evidence of an increase in neuroinflammation and blood coagulation signaling pathways expressed at the neurovascular unit^39, 47, 48^ (Fig. 2). To investigate protein complexes and molecular networks associated with CCM disease-induced neurobehavioral deficits, we conducted a protein profiling study using mass spectrometry on the whole brains of *Pdcd10^BECKO^* mice (Fig 3A). First, proteomic approaches were used to analyze protein abundance in the brains of healthy mice and identify protein changes and relevant pathways across three developmental stages: P15, P40, and P60 (Fig. 3B, C). Proteomics of four biological replicates for each developmental stage yielded 5034 proteins at P15, 5784 proteins at P40, and 5034 proteins at P60 (Supplemental Fig. 3). Gene ontology (GO) analysis of the enriched proteins and functional analysis revealed that the brains of mice at P15 were highly associated with the regulation of actin cytoskeleton, ribosomes, and tight junctions. (Supplemental Fig. 3). In contrast, the brains of mice at P40 and P60 showed high association with proteins linked to several neurological diseases, oxidative phosphorylation, metabolic pathways, and neurotransmitter activity. This suggests that the brains at this developmental stage are more mature in the context of brain functions (Supplemental Fig. 3).

**Figure 3.**
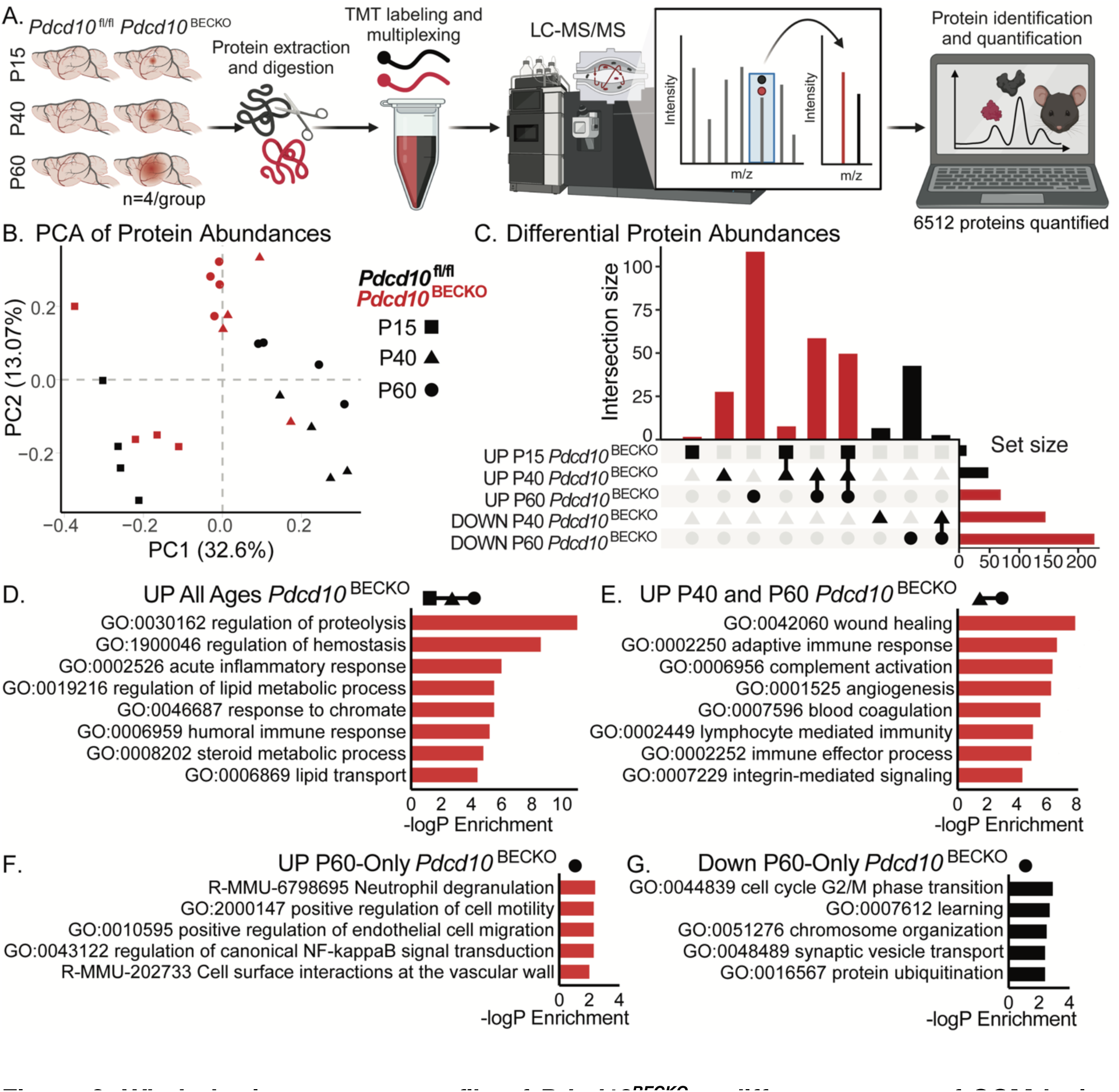
Whole-brain proteome profile of *Pdcd10^BECKO^* at different stages of CCM lesion genesis. A) Schematic diagram of *Pdcd10^fl/fl^* and *Pdcd10^BECKO^* murine brain tissue preparation for quantitative multiplexed proteomics, comparing P15, P40, and P60 aged mice (n=4/group). B) Principal component analysis of all quantified proteins. C) UpSet plot of significantly differentially abundant proteins identified in binary comparisons of *Pdcd10*^BECKO^ versus age-matched *Pdcd10*^fl/fl^ controls by group. Significantly differentially abundant proteins were assessed by a combination of Welch’s t-tests with a p-value of < 0.05 and a π-score of > 1. D-G) Gene ontology analyses of proteins D) up in abundance at all ages in *Pdcd10*^BECKO^: *Pdcd10*^fl/fl^, E) up at ages P40 and P60 in *Pdcd10*^BECKO^: *Pdcd10*^fl/fl^, F) up uniquely at age P60 in *Pdcd10*^BECKO^: *Pdcd10*^fl/fl^, and G) down uniquely at age P60 in *Pdcd10*^BECKO^: *Pdcd10*^fl/fl^.

Moreover, we used integrating multi-omics datasets to identify altered proteins, genes, and pathways that occur during CCM disease (Fig. 3C-G). Notably, GO analysis of altered proteins in CCM disease at all CCM stages revealed functional clusters corresponding to the regulation of proteolysis, hemostasis, and lipid metabolic processes (Fig. 3D). Wound healing, adaptive immune response, complement activation, and neuroinflammation were notably increased at CCM mature lesions (Fig. 3E-G, 4A). Upon further analysis, comparing RNAseq and protein abundance data from *Pdcd10^fl/l/^* mice revealed that protein abundance in healthy brains is largely consistent with the corresponding RNA abundance (Fig. 4B). Our findings reveal that protein changes from whole-brain analysis of CCM disease partially align with the corresponding RNA levels, with many newly differentially abundant proteins uniquely identified in the proteomics study (Fig. 4B, C). However, we discovered proteins previously associated with CCM pathogenesis, such as an increase in GFAP and vimentin as reflective of the increase in CCM reactive astrocytes^4, 39, 48^, hemoglobin protein (HBA-A1, HBB-B1), fibronectin 1^55^, and proteins associated with blood coagulation clothing THBD and TSP1^39, 48^ (Fig. 4C-G). These results strongly suggest that activation of signaling pathways associated with neuroinflammation and thrombosis affects the chronic stage of CCM disease and behavioral outcomes.

**Figure 4.**
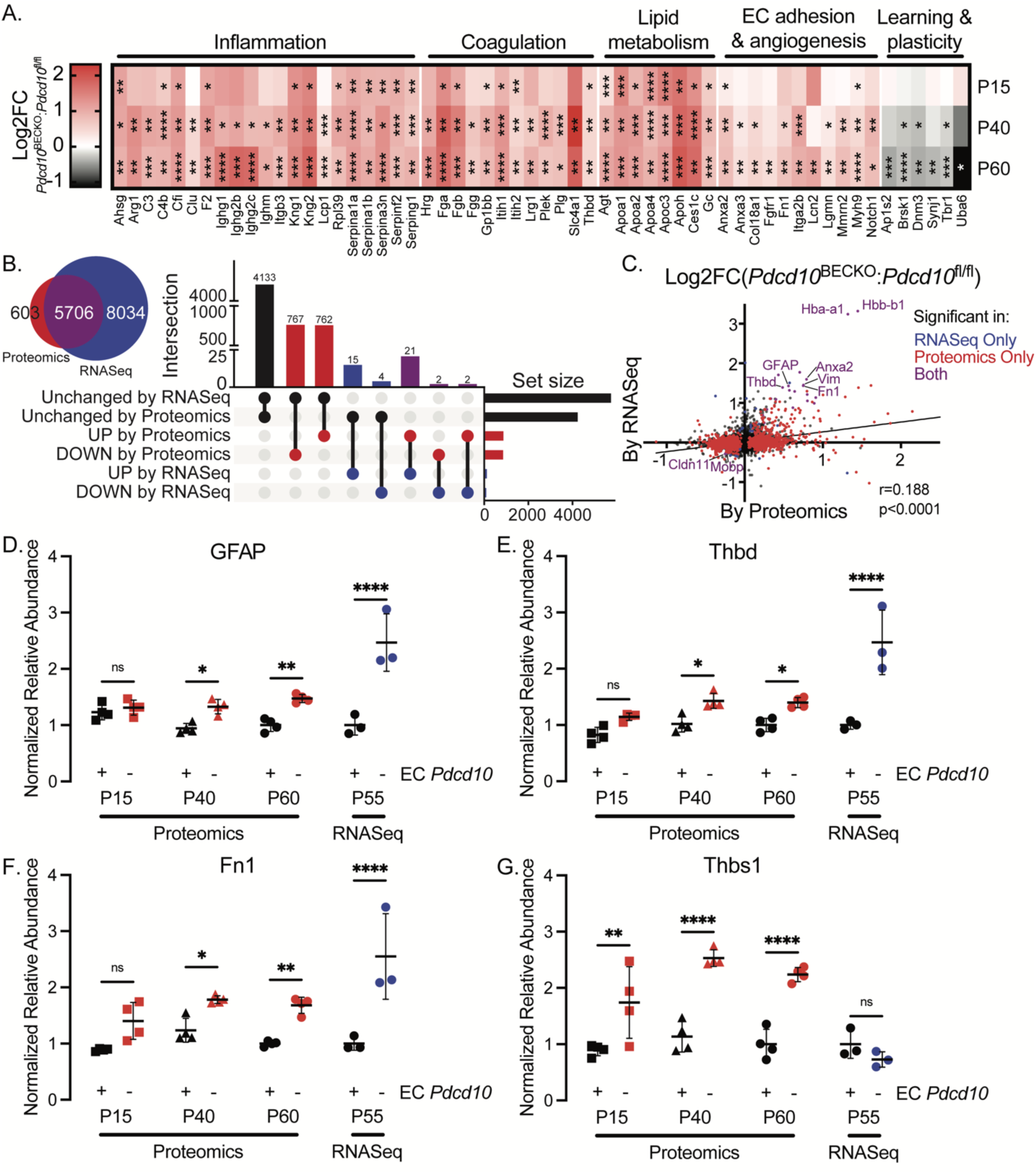
*Pdcd10^BECKO^*whole-brain proteome profiling indicates elevated neuroinflammation and thrombosis, especially by P60. A) Heatmap of differentially abundant proteins quantified in *Pdcd10^BECKO^* murine brain tissue, colored by the fold-change of *Pdcd10*^BECKO^: *Pdcd10*^fl/fl^, organized by ontology. Statistical significance is represented with *, **, ***, and **** indicating p-values of less than 0.05, 0.01, 0.001, and 0.0001, respectively. B) Venn diagram illustrating overlapping detected genes between current proteomics and previous transcriptomics study^48^, and UpSet plot illustrating overlapping and uniquely significant genes in binary comparisons across the study, looking at P60 and P55 mice in the proteomics and transcriptomics studies, respectively. C) Linear correlation between fold-changes of *Pdcd10*^BECKO^: *Pdcd10*^fl/fl^ in present proteomics and previous transcriptomics studies, with significantly different genes in color. D-G) Previously reported markers of CCM are recapitulated by the current proteomics study and show an increase with age. Each graph is normalized to the average P60 of Pdcd10fl/fl, and error bars represent the standard deviation.

## Discussion

Our study revealed that CCM animal models displayed behavioral deficits, including reduced motor coordination and anterograde amnesia. Using histological and proteomic methods, we discovered that the behavioral impairments were associated with the development of CCM lesions showing a neuroinflammatory pattern.

We first noted that lesions in a CNS-specific CCM animal model initially form in the cerebellum^4^ and have minimal impact on the RotaRod performance. This suggests that even when the CCM lesion formation forms extensive dilated vessels in this CCM animal model, it has little effect on motor coordination deficits. Because we observed no significant difference in the performance of *Pdcd10^BECKO^* and control *Pdcd10^fl/fl^* mice on the RotaRod test at P35. Suggesting that rodents may compensate for deformities in brain blood vessels during the development of CCM disease to prevent brain deficits^56^. However, reduced motor coordination was observed in *Pdcd10^BECKO^* mice with neuroinflammatory CCM lesions^4, 39^. Neuroinflammation also drives pathogenic processes, resulting in neuropsychiatric and cognitive effects ^57, 58^. Indeed, we observed that using specific metrics for contextual and cued fear-conditioning tests to assess short- and long-term fear memory indicates that neuroinflammation in *Pdcd10^BECKO^* mice may contribute to significant impairment in both short- and long-term fear memory. A recent study unveiled new insights into the mechanisms of neuroinflammatory CX3CR1-CX3CL1 signaling and the propensity of CCM immunthrombosis. Interactions between CX3CR1 and CX3CL1 modify CCM neuropathology when lesions are accelerated by environmental factors, such as mild hypoxia, and during chronic CCM disease^48^. The authors established CX3CR1 as a genetic marker for patient stratification and a potential predictor of CCM aggressiveness^48^. However, additional studies should conduct experiments to better define the drivers of lesion genesis in comparison to thrombosis drivers. In summary, our analysis suggests that CCM-related neuroinflammation significantly impacts neurobehavioral outcomes. Future preclinical studies in CCM therapeutics^8^ should aim to assess whether they also improve neurobehavioral deficits.

## Authors’ contributions

J.O., B.C., I.J., L.C., and S.A. designed and performed the behavioral experiments, analyzed and interpreted the data, generated the figures, and wrote the manuscript. B.N. assisted with preliminary studies of CCM in the behavior test; E.F-A. and H.G-G. designed, performed, and analyzed histological results in conditional knockout mice and helped to review and edit the manuscript. L-A.R. and D.J.G. designed, performed, and analyzed data from proteomic experiments, generated the figures, and helped to review and edit the manuscript. M.A.L-R. conceived the project, designed the overall study, analyzed and interpreted data, and wrote the manuscript.

## Acknowledgments

The authors thanks Connie Lee, Jianbo Hu, and Amy Akers from Alliance to Cure Cavernous Malformation for helpful discussion; Cassandra Bui, for technical assistance; Jennifer Santini and Marcy Erb for microscopy technical assistance; *Slco1c1-CreERT2 mice* were the generous gift of Markus Schwaninger (University of Lübeck); *Pdcd10^fl/fl^*mice were the generous gift of Wang Min (Yale University).

## Supplemental figures

**Supplemental Figure 1.**
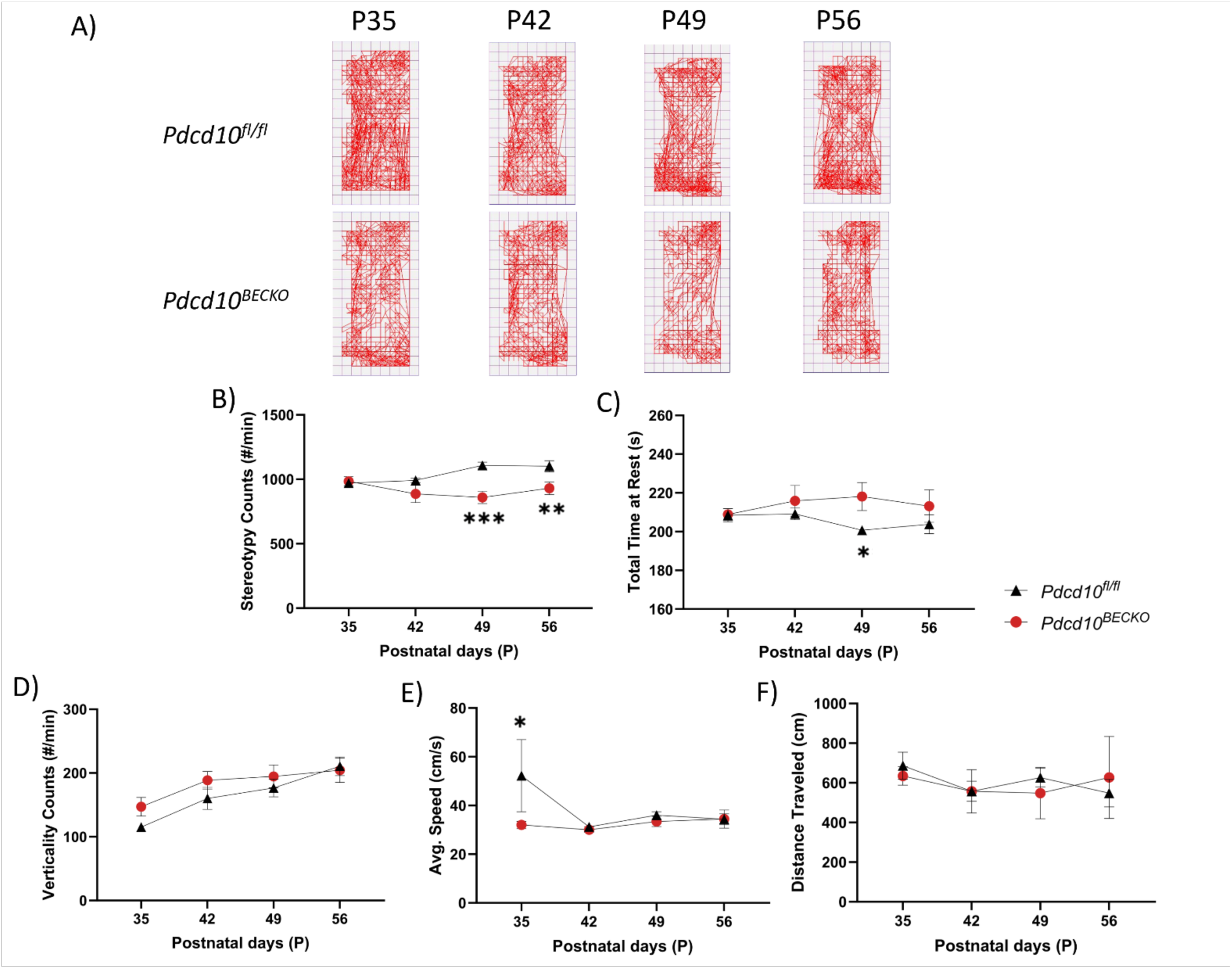
P*d*cd10BECKO mice Open Field Test. A) Representative track plot reports recorded during the 30 min test sessions (Med Associates Inc.). B) *Pdcd10^BECKO^* male and female mice show decrease in stereotypy after P42 (n = 8 *Pdcd10^fl/fl^* and 6 *Pdcd10^BECKO^*). C) *Pdcd10^BECKO^* male and female mice show increased resting time at P49. D) No significant differences were found in verticality between groups. E) *Pdcd10^BECKO^* male and female mice show a decrease in average movement speed at P35. F) No significant differences were found between groups in distance traveled. Data are presented as a mean ± SEM. \**P* < 0.05, \*\**P* < 0.01, \*\*\**P* < 0.001.

**Supplemental Figure 2.**
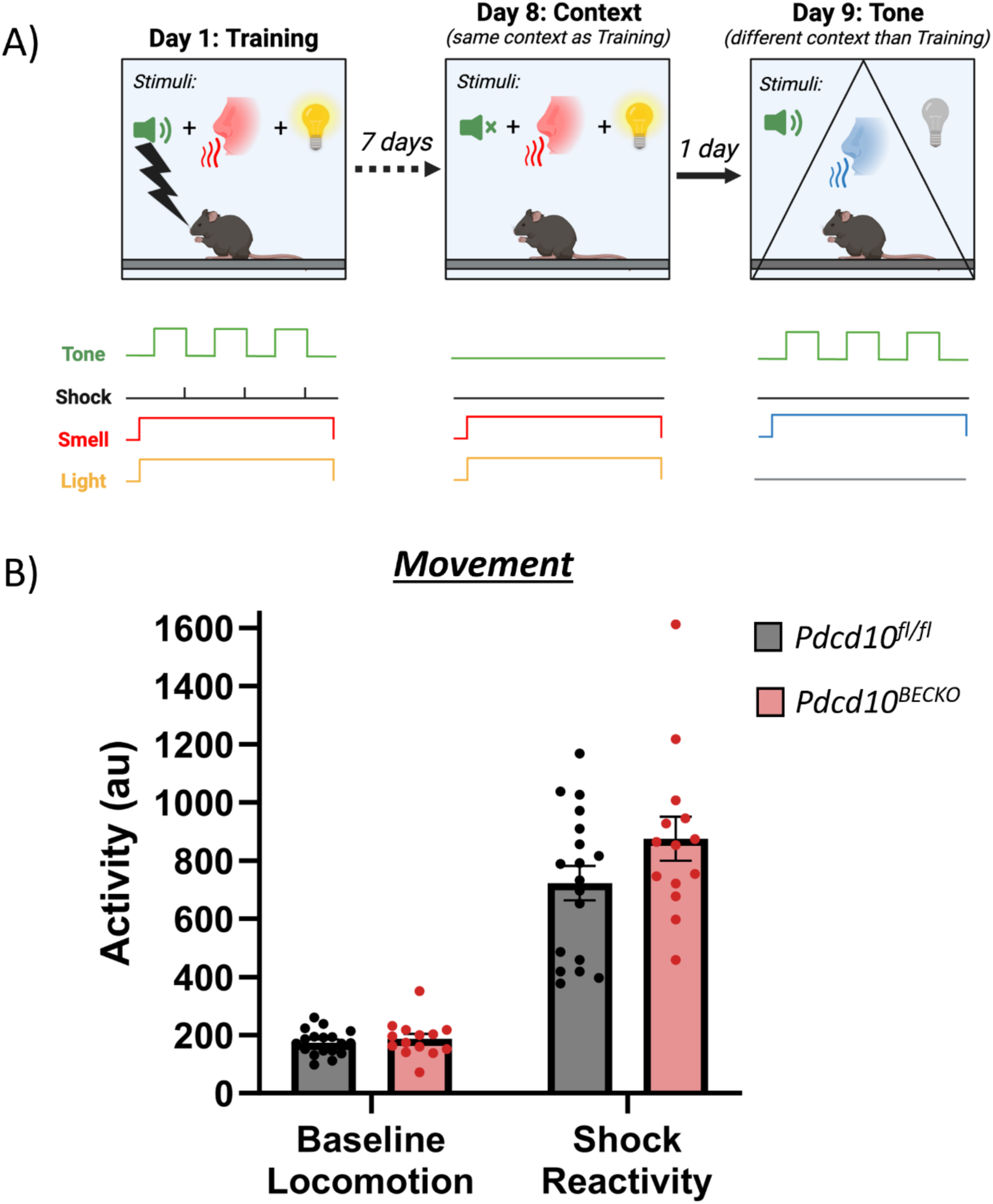
P*d*cd10BECKO mice Fear Conditioning Assessment. A) Experimental setup for Pavlovian fear conditioning in mice. On Day 1 (Training), mice were placed in a conditioning box scented with isopropyl alcohol, with the lights turned on. The mice experienced a 2-minute acclimation period, followed by three consecutive trials of a 30-second auditory tone co-terminating with a mild footshock, with intervals of 30 seconds between trials. Post-trial, mice remained in the box for an additional 5 minutes to assess short-term memory recall. On Day 8 (Context), mice were reintroduced to the identical box environment from the training phase (minus the tone and shock) to evaluate recognition of the context and long-term context memory associated with fear. Finally, on Day 9 (Tone), mice were placed in a novel environment with distinct walls, floors, and scent, without lighting. The altered environment was coupled with three repetitions of the 30-second tone (with no footshock) to examine long-term memory recall of the tone. The experiment measures the mice’s memory and fear responses to the conditioning stimuli over time. B) Baseline locomotion and shock reactivity during the 2min training baseline period and 2s footshock. While group differences were observed in short- and long-term fear memory, no differences were observed in baseline locomotion (n = 18 *Pdcd10^fl/fl^*and 14 *Pdcd10^BECKO^*) or reactivity to the footshock.

**Supplemental Figure 3.**
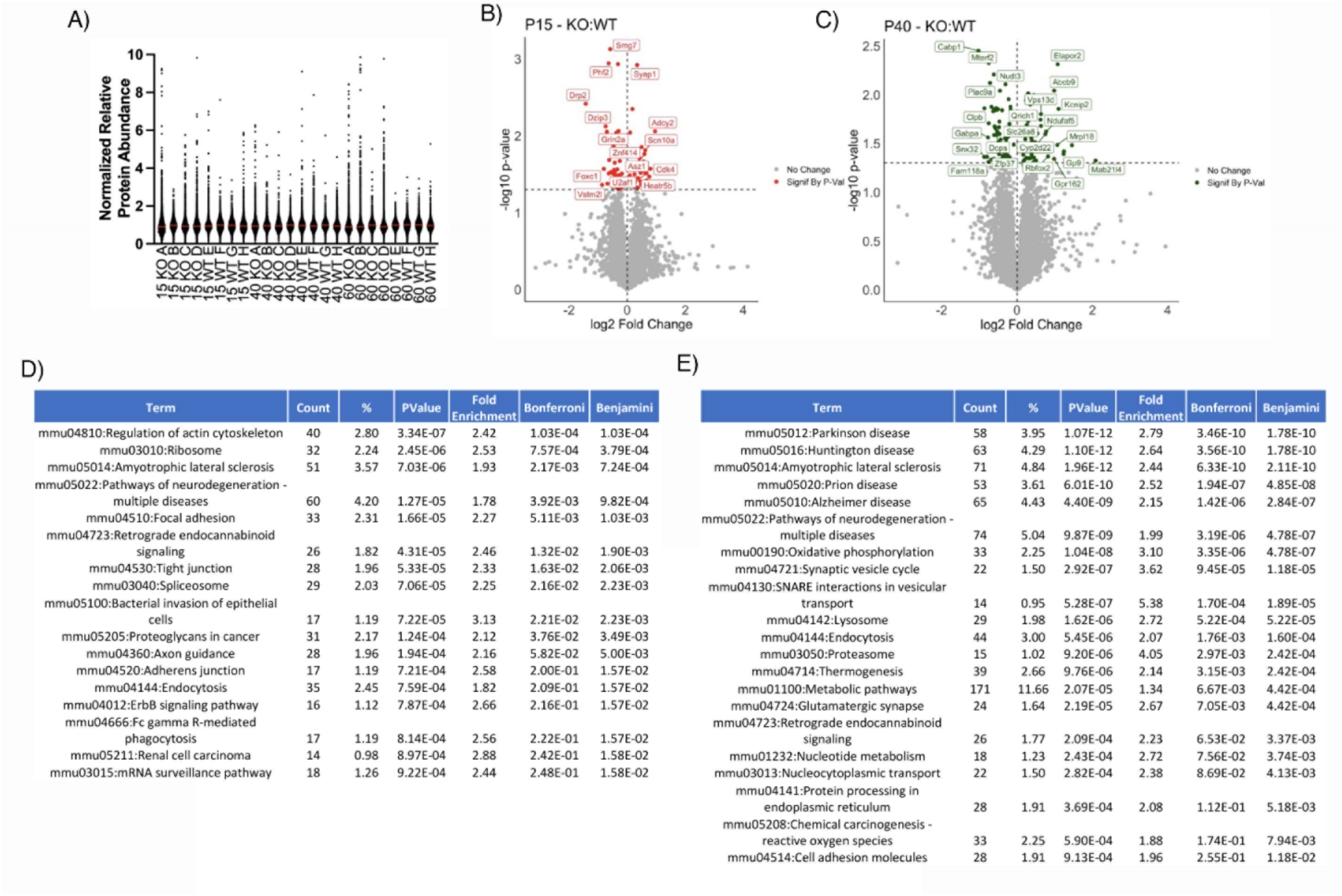
P*d*cd10BECKO whole-brain proteome profiling. A) The graph depicts the distribution of peptide abundance in *Pdcd10^fl/fl^* and *Pdcd10^BECKO^*murine brain tissue at various developmental stages (P15, P40, and P60) as indicated. B-C) Volcano plot of peptides quantified from *Pdcd10^fl/fl^* and *Pdcd10^BECKO^* murine brain tissue at P15 (*B*) and P40 (*C*). The −log10-transformed *p* values associated with individual peptides are plotted against the log2-transformed fold change in abundance between *Pdcd10^fl/fl^* and *Pdcd10^BECKO^* brains. Color intensities depict peptides with significantly higher (right) or lower (left) levels in *Pdcd10^BECKO^* compared to *Pdcd10^fl/fl^*. D) Mass spectrometry analysis of whole-brain samples was used to analyze protein levels in the brains of P15 *Pdcd10^fl/fl^* mice. E) D) Mass spectrometry analysis of whole-brain samples was used to analyze protein levels in the brains of P40 *Pdcd10^fl/fl^* mice.

